# 3D bioprinting of co-cultured osteogenic spheroids for bone tissue fabrication

**DOI:** 10.1101/2020.06.16.155143

**Authors:** Dong Nyoung Heo, Bugra Ayan, Madhuri Dey, Dishary Banerjee, Hwabok Wee, Gregory S. Lewis, Ibrahim T. Ozbolat

## Abstract

Conventional top-down approaches in tissue engineering involving cell seeding on scaffolds have been widely used in bone engineering applications. However, scaffold-based bone tissue constructs have had limited clinical translation due to constrains in supporting scaffolds, minimal flexibility in tuning scaffold degradation, and low achievable cell seeding density as compared with native bone tissue. Here, we demonstrate a pragmatic and scalable bottom-up method, inspired from embryonic developmental biology, to build three-dimensional (3D) scaffold-free constructs using spheroids as building blocks. Human umbilical vein endothelial cells (HUVECs) were introduced to human mesenchymal stem cells (hMSCs) (hMSC/HUVEC) and spheroids were fabricated by an aggregate culture system. Bone tissue was generated by induction of osteogenic differentiation in hMSC/HUVEC spheroids for 10 days, with enhanced osteogenic differentiation and cell viability in the core of the spheroids compared to hMSC-only spheroids. Aspiration-assisted bioprinting (AAB) is a new bioprinting technique which allows precise positioning of spheroids (11% with respect to the spheroid diameter) by employing aspiration to lift individual spheroids and bioprint them onto a hydrogel. AAB facilitated bioprinting of scaffold-free bone tissue constructs using the pre-differentiated hMSC/HUVEC spheroids. These constructs demonstrated negligible changes in their shape for two days after bioprinting owing to the reduced proliferative potential of differentiated stem cells. Bioprinted bone tissues showed interconnectivity with actin-filament formation and high expression of osteogenic and endothelial-specific gene factors. This study thus presents a viable approach for 3D bioprinting of complex-shaped geometries using spheroids as building blocks, which can be used for various applications including but not limited to, tissue engineering, organ-on-a-chip and microfluidic devices, drug screening and, disease modeling.

## 1. Introduction

Bone is a highly vascularized dynamic tissue, observed in a variety of shapes and sizes in the body. The skeleton serves crucial roles such as supporting the framework for the body and protecting vital organs [1,2]. Large bone defects cannot self-regenerate regenerate, and can be due to trauma, fracture nonunion, infection, bone tumor resections, and removal/revision of joint replacements and other implant [3–5]. There is thus a substantial demand for engineered constructs to restore and regenerate the diseased/excised part of the bone tissue [4,6,7]. To-date, significant progress has been made using three-dimensional (3D) bioprinting of living cells and tissues to recapitulate the bone tissue [8]. Most of these bioprinting approaches rely on scaffold-based techniques where biodegradable hydrogels are seeded or combined with mature osteoblasts or osteogenically-committed stem-cells [6,9,10]. These scaffolds can provide mechanical support, serve as a template for cell attachment and facilitate a conducive environment for cellular activities [3,6]. However, degradation of scaffolds, limited cell density compared to native-tissues and limited communication among cells are some of the major drawbacks in scaffold-based approaches [11].

Scaffold-free bone tissue engineering pose a promising alternative to 3D bioprinting approaches [12]. Stem-cell derived aggregates, in the form of spheroids, are considered as building blocks for scaffold-free bioprinting and mimic the complex morphology and physiology of native tissue by inducing cross-talk among cells and cell-extracellular cell matrix (ECM) interactions [13,14]. In addition, osteogenic differentiation is observed to increase due to stronger integrin-ECM interaction caused by the presence of both stroma and structure within spheroids [15]. However, non-uniform bone tissue regeneration owing to the non-homogeneous oxygen diffusion across the entire spheroid domain, especially to the core of spheroids [15–17] forms the major roadblock in successful usage of spheroids in 3D bioprinting applications. 2D co-culture systems with mature osteoblasts or stem cell derived osteogenic progenitor cells with cells from endothelial lineage have shown potential to address this limitation by enhanced secretion of endothelial cell-mediated paracrine factors secretion under hypoxia [18]. In this study, we intended to exploit this potential of endothelial cells and investigate their role in 3D on osteogenic differentiation. We achieved this by the co-culture of a minimal number of human umbilical vein endothelial cells (HUVEC) in human mesenchymal stem cells (hMSC) spheroids, and by induction of osteogenesis with bone regeneration across the entire domain of spheroids.

Spheroids have been utilized in successful biofabrication of functional bone tissue substitutes [19–22]. Although several approaches have been presented in the literature for bioprinting of spheroids, most of these suffer from poor spatial control in 3D, significant damages to spheroids with loss of viability and structural integrity, poor repeatability of the process while using spheroids that are non-uniform in size, inability to form complex 3D shapes, inability to maintain designed shape due to cell-mediated compaction post-bioprinting, and lack of scalability for translation from bench to bedside [19–21]. Aspiration-assisted bioprinting (AAB), a newly developed technique by our group, bioprints spheroids in 3D space using the power of aspiration forces. AAB leverages the fact that spheroids can be formed from diverse cell types at high densities [22]. When using human stem cells, these spheroids can recapitulate aspects of embryonic development to self-assemble [21] into various organs. Using this technique, in this study, we have demonstrated that spheroids with viscoelastic properties can be lifted by aspiration forces, and then positioned precisely (11% with respect to the spheroid diameter) into a hydrogel substrate, circumventing the limitations of the other spheroid bioprinting techniques available.

Here, co-culture of hMSCs and HUVEC spheroids were utilized to enhance cellular function, osteogenic differentiation, calcium deposition, and cell viability. After biofabrication of spheroids, two different strategies were demonstrated for the bioprinting of bone tissue using hMSC/HUVEC and pre-differentiated hMSC/HUVEC spheroids. Compaction of spheroids leading to significant changes in geometry of tissue constructs compared to the desired has been a common issue post bioprinting, and hence, we utilize 3D bioprinting of pre-differentiated hMSC/HUVEC spheroid to minimize shape changes. Post-bioprinting, we showed that we were able to control the shape of the bioprinted construct, using pre-differentiated hMSC/HUVEC spheroids. In this study, thus, we attempted to address major limitations of spheroid bioprinting by bioprinting spheroids using AAB, fabricating complex-shaped bone tissue constructs, engineering the spheroids in a way to induce osteogenesis across the entire spheroid domain, and controlling osteogenic induction timelines before and after bioprinting in order to reduce spheroid compaction and increase retention of geometry of the tissue constructs.

## 2. Experiment Section

### 2.1. Cell culture

HMSCs (Lonza, Walkersville, MD) and HUVECs (Lonza) were used to fabricate of 3D cellular spheroids. hMSCs were cultured in all-in-one ready-to-use hMSC growth medium (Cell Applications, INC., San Diego, CA). HUVECs or tdTomato^+^ HUVECs were cultured in MCDB 131 medium (Corning, New York, NY) supplemented with 10% fetal bovine serum (Corning, New York, NY), 2 mM glutamine (Thermo Fisher Scientific, Waltham, MA), 1% penicillin/streptomycin (Corning, New York, NY), 4.5 ug/mL bovine brain extract (Lonza, Walkersville, MD), 10 unit/mL heparin (Sigma-Aldrich, St. Louis, MO), and 1.86 mg/mL endothelial cell growth supplement (Sigma-Aldrich, St. Louis, MO). Cells passages from three through seven were used for both hMSCs and HUVECs. The cells were expanded at 37 °C with 5% CO_2_ in a humidified sterile incubator.

### 2.2. Spheroid fabrication

hMSCs and HUVECs were harvested with trypsin and collected by centrifugation at 1600 rpm for 5 min for the fabrication of the spheroids. hMSCs were reconstituted to 2.5 × 10^5^ cells per ml, with various mixing ratios of HUVECs (8, 15, and 25 %), referred as 92:8, 85:15 and 75:25 hMSC/HUVEC respectively. 200 μl of cell suspension was seeded into each well of 96-well plates with U-bottom, cell-repellent surface (Greiner bio-one, Frickenhausen, Germany) to achieve ∼50,000 cells/spheroid. Cells were maintained in hybrid growth media (75% MSC and 25% HUVEC growth media) to allow compaction at 37 °C. HMSC-only spheroids were fabricated similarly and used as control to understand the functionality of HUVECs in the spheroids. Osteogenic differentiation was induced in hMSC-only spheroids and hMSC/HUVEC spheroids by cultured in human osteogenic differentiation medium (Cell Applications, INC., San Diego, CA) for 10 days. The cell medium was changed every three days.

### 2.3. Characterization of spheroids

The diameters of hMSC-only and hMSC/HUVEC spheroids were imaged and further quantified using EVOS FL cell imaging system (Thermo Fisher Scientific, Waltham, MA) and ImageJ (NIH Freeware) respectively from each of days, 1 to 5 post-seeding. Cell viability was confirmed using a calcein-AM/ethidium homodimer-1 (EthD-1) LIVE/DEAD assay kit (Invitrogen, Carlsbad, CA) [22]. Briefly, five days after incubation, spheroids were washed with Dulbecco’s phosphate-buffered saline (DPBS), stained with 2 μM calcein AM and 4 μM EthD-1 solutions for 40 min, and then imaged as described before.

To evaluate the cellular organization of hMSC-only and hMSC/HUVEC spheroids, spheroids were fixed in 10 % formalin, embedded in paraffin using a Leica TP 1020 automatic tissue processor (Leica, Wetzlar, Germany) and sectioned with 10-μm thickness. Cross-sectioned samples were stained with hematoxylin and eosin (H&E) using a Leica Autostainer XL (Leica, Wetzlar, Germany) and then imaged using optical microscopy. In addition, the cytoskeletal organization was confirmed with actin staining. Cross-sectioned samples were permeabilized in 0.1% Triton X-100 for 30 min, blocked with 2.5% normal goat serum (NGS) for 60 min at room temperature and incubated with ActinGreen™ 488 (1:1000 in 2.5% NGS) and 4’,6-Diamidino-2-Phenylindole (DAPI; 1:1000 in 2.5% NGS) for 60 min. Then, the stained samples were washed thrice with DPBS and observed under an optical microscope.

### 2.4. Surface tension measurements

The surface tension of hMSC-only and hMSC/HUVEC spheroids was measured using a micropipette aspiration technique, as previously described [23]. The customized micropipettes were prepared from borosilicate Pasteur pipettes (vWR, 14673-043, Radnor, PA) on a P1000 Flaming/Brown micropipette puller (Sutter Instrument, Novato, CA). Parameters for heat, pull, velocity, time, and pressure were set at 575, 0, 5, 100, and 50, respectively according to our recent study [22]. All spheroid types were aspirated by a customized glass pipette and monitored via an STC-MC33USB monochromatic camera (Sentech, Japan) equipped with 1-61448 and 1-61449 adaptor tubes (Navitar, Rochester, NY).

### 2.5 Quantitative real-time polymerase chain reaction (real-time PCR)

Real-time polymerase chain reaction (RT-qPCR) was performed after five days in growth media, and an additional 10 days in osteogenic media to evaluate the gene expression profiles of hMSC-only and hMSC/HUVEC spheroids. The total RNA of hMSC-only and hMSC/HUVEC spheroids was isolated using a RNeasy Plus Mini Kit (Qiagen, Germantown, MD) according to the manufacturer’s instructions and quantified using a Nanodrop ND-1000 Spectrophotomer (Thermo Scientific, Wilmington, DE). The primers of the measured mRNA genes were as follows: OCT4 (forward, GCA GCG ACT ATG CAC AAC GA and reverse, CCA GAG TGG TGA CGG AGA CA), PECAM-1 (forward, TAA TAC AAC ATC CAC GAG GGT CC and reverse, ACA AAA TTG CTT GCT AAA GAA GTG G), HIF1A (forward, CCA GTT AGG TTC CTT CGA TCA GT and reverse, TTT GAG GAC TTG CGC TTT CA), COL6A1 (forward, CCT GGA GGG CTA CAA GGA A and reverse, GTG CTT GGC CTC GTT CAC), COL1 (forward, ATG ACT ATG AGT ATG GGG AAG CA and reverse, TGG GTC CCT CTG TTA CAC TTT), ALP (forward, AGC TGA ACA GGA ACA ACG TGA and reverse, CTT CAT GGT GCC CGT GGT C), BSP (forward, AAC GAA GAA AGC GAA GCA GAA and reverse, TCT GCC TCT GTG CTG TTG). RT-qPCR was analyzed using SsoFast™ EvaGreen^®^ Supermix (Bio-Rad, Hercules, CA) and all values were normalized by a house-keeping gene GAPDH. Threshold cycle values were calculated using a comparative cycle threshold method. The fold-change of hMSC-only was set at 1-fold, and the ratio of the normalized fold-change was calculated based on the standard condition.

### 2.6 3D bioprinting of spheroids

AAB system was used to bioprint hMSC-only and hMSC/HUVEC spheroids as previously described [22]. AAB system was equipped with microvalves (INKX0517500A, Lee Company, Bashville, TN) with 250-μm nozzles (INZA3100914K, Lee Company, Bashville, TN) and a portable ultrasonic humidifier (CZHD20, Comfort Zone, China). 1g of sodium alginate (Sigma-Aldrich, UK) was dissolved in 100 ml deionized (DI) water to prepare 1% (w/v) sodium alginate as a sacrificial material. In order to generate alginate droplets using the microvalve dispenser, dwell time of 700 μs was utilized with positive back pressure of ∼103 kPa. Calcium chloride (CaCl_2_) solution was prepared by dissolving 4% (w/v) (CaCl_2_ (Sigma-Aldrich, St. Louis, MO) in DI water. Sodium alginate was dispensed on the glass substrate using microvalves, then the aerosol form of CaCl_2_ was utilized to crosslink sodium alginate partially. Spheroids were collected into 1.5 ml conical tubes and transferred to the bioprinting platform. Afterwards, the top portion of the conical tubes were cut by a scissor. A customized glass pipette (∼80 μm in diameter) was dipped into a conical tube and air pressure of 25 mmHg was applied to lift spheroids. Critical lifting pressure was determined theoretically from the mean radius of spheroids, surface tension coefficient of cell media and air interface, and the dynamic contact angle at three phases (air, spheroid, and cell media) [22] to pick spheroids. hMSC-only and hMSC/HUVEC spheroids were lifted from cell media with backpressure of 43 and 52 mmHg, respectively. After the fabrication of complex-shaped arrangements, the bioprinted constructs were overlaid with alginate and crosslinked with the aerosol form. The whole construct was then transferred to a Petri dish and cultured with tissue-specific cell media. 4% (w/v) sodium citrate solution was added and agitated gently to remove alginate.

### 2.7 Characterization of osteogenic differentiation by immunocytochemistry and alizarin red S staining

To visualize the morphologies of osteogenically differentiating spheroids, spheroids cultured in osteogenic differentiation medium were stained with runt-related transcription factor 2 (RUNX2) and CD31. After osteogenic differentiation for 10 days, hMSC-only and hMSC/HUVEC spheroids were cross-sectioned, permeabilized in 0.2% Triton X-100 for 30 min and blocked with 2.5% NGS for 1 h. Then, the samples were incubated with mouse anti-RUNX2 primary antibody (1:100 in 2.5% NGS) and rabbit anti-CD31 primary antibody (1:100 in 2.5% NGS) for 60 min at room temperature, washed three times with DPBS, and incubated with goat anti-mouse Alexa Fluor 488 secondary antibody (1:250 in 2.5% NGS), goat anti-rabbit Alexa Fluor 647 secondary antibody (1:250 in 2.5% NGS), rhodamine-phalloidin (1:250 in 2.5% NGS) and DAPI (1:1000 in 2.5% NGS) for 60 min at 4 °C. The stained samples were then washed three times with DPBS and imaged using an Olympus FV10i-LIV Confocal Laser Scanning Microscope (Olympus America Inc., Melville, NY). The calcium deposition was visualized by staining cross-sectioned slides with a 2 % alizarin red S staining solution for 10 min at room temperature. Stained samples were washed three times with DI water and imaged using optical microscopy.

### 2.8 Micro-computed tomography (μCT) measurements

μCT scanner (VivaCT 40, Scanco Medical, Switzerland) was used with 10.5 μm voxel resolution, 55 kV energy, 145 μA intensity, 21.5mm diameter field-of-view, and 300 ms integration time to evaluate the mineralization of spheroids in bioprinted tissues. The samples were placed inside the μCT scanner and scanned. DICOM files were processed in Avizo software (FEI Company, Hillsboro, OR). A hydroxyapatite (HA) phantom (Micro-CT HA, QRM, Germany) was included in each scan to generate a standard curve to convert Hounsfield units to mgHA/ccm [24]. Images were processed with a Gaussian smoothing filter (sigma 0.9) to reduce noise, and a threshold of 200 mgHA/ccm was used to visualize mineralized spheroids and quantify mineralized volume.

### 2.9 Statistical analysis

All values are presented as mean (±) standard deviation. Multiple comparisons were analyzed using a one-way analysis of variance (ANOVA) followed by Tukey’s multiple comparison test. The differences with p-values (\**p* < 0.05, \*\**p* < 0.01) were considered statistically significant. All statistical analysis was performed by Statistical Product and Service Solutions software (SPSS, IBM, Armonk, NY).

## 3. Results and Discussion

### 3.1 Fabrication and characterization of hMSC-only and hMSC/HUVEC spheroids

96-well plates with cell-repellent surface were used to form spheroids composed of hMSC-only and hMSC/HUVEC as described in our previous work [13]. As shown in **Figure 1A**, optical microscopy images demonstrated that cell suspension with a density of 50,000 cells per well was undergoing formation of a 3D cellular network, where cells coalesced into a single spheroid along the bottom surface of the well plate. Once aggregation was completed, spheroids slightly decreased in size, but progressively compacted over the course of culture period (**Figure 1A**). All spheroids demonstrated a spherical and uniform shape at day 5 of culture. The mean diameter of hMSC-only, and 92:8, 85:15, and 75:25 hMSC/HUVEC spheroids was determined to be ∼ 694, ∼655, ∼649, and ∼592 μm, respectively (**Figure 1B**). Cell viability of spheroids at day 5 using LIVE/DEAD staining depicted ∼86, and ∼92% for hMSC-only, and all hMSC/HUVEC spheroids respectively (**Figure 1C**). The cell viability of all the groups introduced with HUVECs was found to be significantly higher than that of hMSC-only group, which is in accordance to previous 2D co-culture reports [25]. In particular, cell viability was uniform across the entire spheroid domain for HUVEC-involved groups. H&E staining images (**Figure 1A**) of cross-sections of spheroids confirmed evenly distributed and well-connected cells with no significant difference in cell viability (**Figure 1C**) when comparing the number of HUVECs in hMSC spheroids. Also, immunofluorescent staining of F-actin corroborated the results from the H&E staining. hMSC/HUVEC spheroids exhibited denser and more uniform cytoskeletal organization as compared to hMSC-only spheroids, which exhibited lesser cellular density and empty space at the core. Moreover, hMSC/HUVEC spheroids had significantly higher RNA amount. As shown in **Figure 1D**, the RNA amount for hMSC-only, 92:8, 85:15, and 75:25 hMSC/HUVEC spheroids was ∼79, 126, 162, and 196 ng/spheroid, respectively. Interestingly, the RNA amount of hMSC/HUVEC spheroids decreased with increase in HUVEC concentration. This showed that larger cellular spheroids needed to generate a higher amount of RNA for the maintenance of cellular activities [26]. Thus, the dead empty core with reduced cell viability and RNA amount in hMSC-only spheroid are a direct evidence to the role of HUVECs in enhancing the crosstalk amongst hMSCs in the spheroids. Moreover, visibly more compaction, also corroborated by the diameters of the spheroids, in the hMSC/HUVEC spheroids improved the diffusion of the nutrients into the core of the spheroids and facilitated the removal of waste products, enhancing the cell viability.

**Figure 1.**
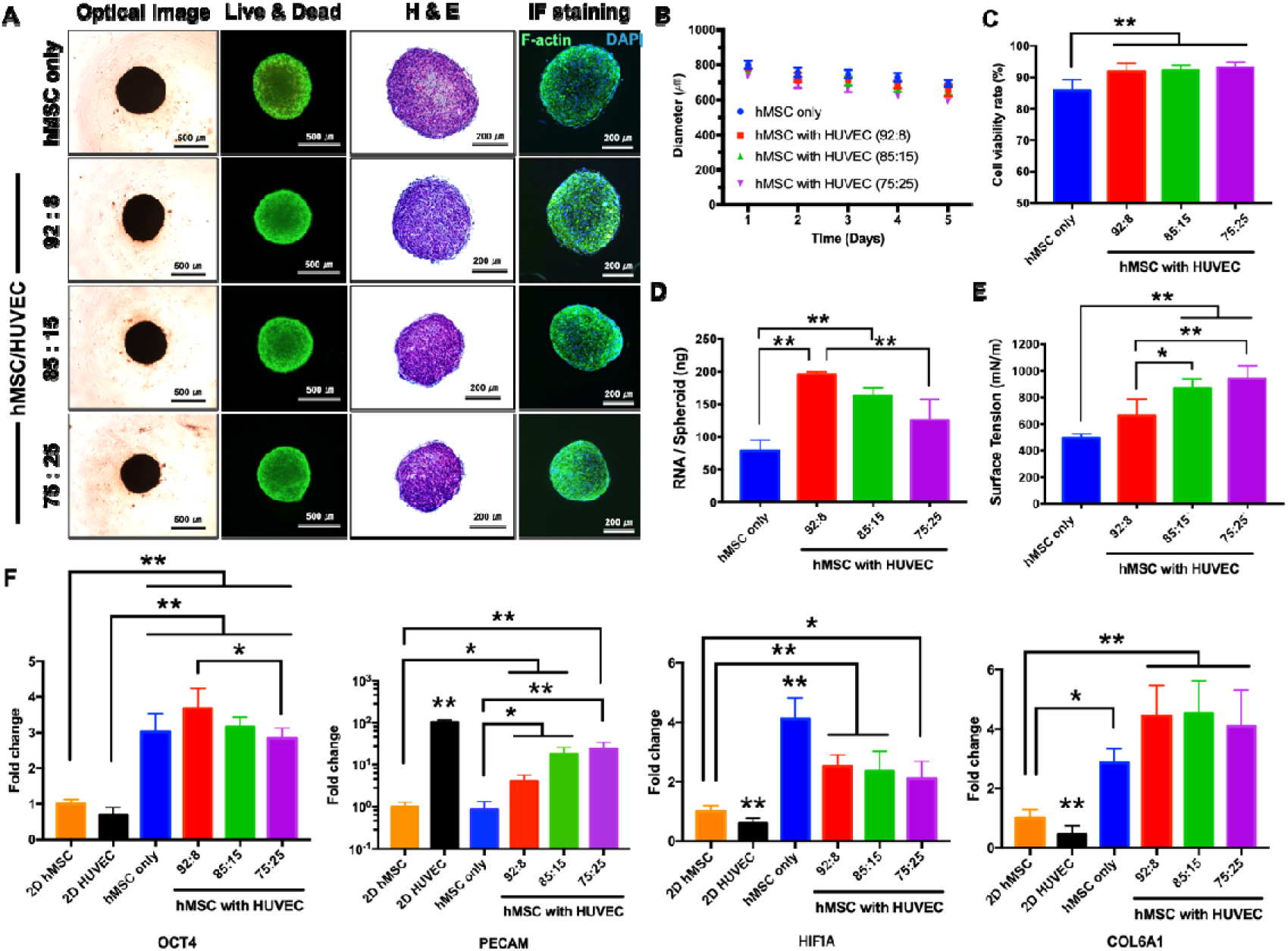
Formation of 3D spheroids and their characterization five days after incubation: **(A)** optical, LIVE/DEAD assay, histology, and F-actin staining images of hMSC-only and hMSC/HUVEC spheroids, **(B)** compaction of the spheroids over time, determined from the reduction in their spheroid diameter. **(C)** cell viability, quantified from LIVE/DEAD assay **(D)** quantification of RNA concentration using Nanodrop **(E)** surface tension **(F)** gene expression measurements for pluripotent stem cell (OCT4), endothelial cell (PECAM), hypoxia (H1F1A), and ECM (COL6A1) related markers in 2D hMSC, hMSC-only and hMSC/HUVEC spheroids; *p < 0.05 and **p < 0.01. We showed that introduction of only 8% HUVECs into the hMSCs is sufficient enough to enhance cell viability at the core of the spheroid with improved mechanical properties.

The surface tension of spheroids was measured as 495.8 ± 30.2 mN/m for hMSC-only spheroids, 661.9 ± 127.1 mN/m, 865.7 ± 73.8 mN/m, and 940.6 ± 98.8 mN/m for 92:8, 85:15, and 75:25 hMSC/HUVEC spheroids respectively (**Figure 1E**). Notably, all hMSC/HUVEC groups showed increase in surface tension as compared to that of hMSC-only spheroid. The 75:25 group showed the highest surface tension amongst all groups, attributed to decrease in diameter of hMSC/HUVEC spheroid with increase in HUVEC concentration, as discussed earlier (**Figure 1B**). This shows that the compaction of the spheroids, depicted by the decrease in diameter, is a direct influencer of the surface tension of the spheroids. The increased compaction demonstrates that the hMSC/HUVEC spheroids are more in the viscoelastic spectrum, i.e. more elastic (solid) than viscous (liquid), and hence possess enhanced mechanical properties compared with hMSC-only spheroids.

In order to quantify the gene expression profiles of hMSC-only and hMSC/HUVEC spheroids, mRNA expression of OCT4, PECAM-1, HIF1A, and COL6A1 genes was investigated by RT-qPCR. As shown in **Figure 1F**, the mRNA expression of OCT4, represented as pluripotency-associated gene [27], significantly upregulated for hMSC-only and hMSC/HUVEC spheroids compared to conventional 2D culture condition of hMSC (2D hMSC) and HUVEC (2D HUVEC) after five days of culture in growth media. Among all groups, 92:8 hMSC/HUVEC spheroids showed the highest expression level of OCT4 compared to that for other groups. Additionally, all hMSC/HUVEC spheroids exhibited higher PECAM-1 gene expression compared to 2D hMSC and hMSC-only spheroids. PECAM-1 expression level was significantly increased with increase in HUVEC concentration and 75:25 group exhibited the highest PECAM-1 expression. On the other hand, the expression of HIF1A, which is a hypoxia-responsive transcription factor [28], was significantly decreased with increase in HUVEC concentration. The highest expression level of HIF1A was observed in hMSC-only spheroid. COL6A1 is a major structural component of microfibrils in connective tissues and interact with other ECM components [29]. COL6A1 expression of hMSC-only and hMSC/HUVEC spheroids was significantly higher than that of 2D hMSC and 2D HUVEC. Overall, all hMSC/HUVEC groups showed increased mRNA expression of OCT4, PECAM, and COL6A1 and decreased HIF1A expression compared to hMSC-only spheroids.

In summary, 92:8 hMSC/HUVEC spheroids showed higher cell viability and mechanical properties (surface tension) compared to hMSC-only spheroids and demonstrated highest RNA content and pluripotency potential. Based on these results, we showed that introduction of only 8% HUVECs into the hMSCs is sufficient enough to enhance spheroid core cell viability with improved mechanical properties. Therefore, to accomplish our goal of building bone tissue and exploring the role of HUVECs in hMSCs spheroids on shape preservation of 3D constructs, we utilized 92:8 hMSC/HUVEC spheroids for the rest of the study, referred as hMSC/HUVEC spheroid henceforth.

### 3.2 Bioprinting of hMSC/HUVEC spheroids via AAB

In our previous study, we developed an AAB platform for biofabrication of 3D constructs with complex geometries [22]. This AAB system can precisely bioprint spheroids with a diverse range of sizes (diameter <80 μm − >800 μm) by picking up spheroids individually and placing them onto desired locations on hydrogel-coated substrates. In this study, alginate was printed onto glass substrates using a micro-valve dispenser, and partially cross-linked using aerosol of CaCl_2_. hMSC/HUVEC spheroids were picked by a glass pipette and bioprinted at desired locations onto the gel, according to previously determined patterns (**Figures 2A–C, Supplementary Video 1)**. This procedure was repeated for each of the spheroids and after completion, the spheroids were stacked up in three layers to fabricate each of the 3D complex-shaped constructs in mm scale, resembling a pyramid (**Supplementary Video 1)**, hexagon, and diamond topology. Owing to the inherent adhesive nature of hMSC/HUVEC spheroids ^21,^ [22], they were well stacked up on the gel substrate and maintained the 3D complex topologies without structural collapse.

**Figure 2.**
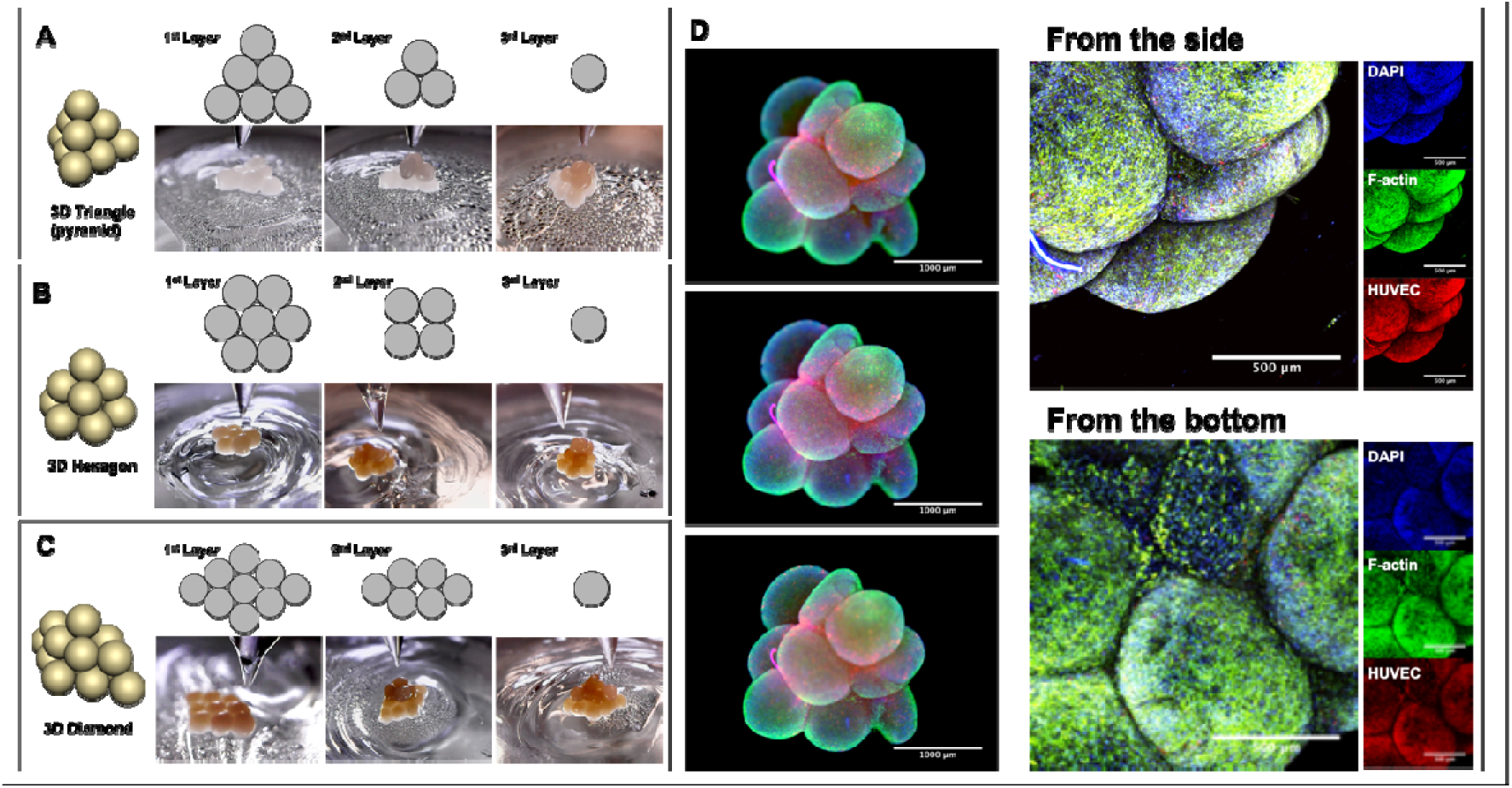
3D Bioprinting of hMSC/HUVEC spheroids via Aspiration-assisted Bioprinting. Schematic diagrams and real-time photographs of a 3D pyramid **(A)**, 3D hexagon **(B)**, and 3D diamond **(C)** topographies using spheroids as building blocks. **(D)** Fluorescent images of bioprinted pyramid of hMSC/ tdTomato^+^ HUVEC spheroids, which were stained with DAPI and F-actin. Confocal images of pyramid constructs (side and bottom views).

After bioprinting, the constructs were overlaid with alginate using the micro-valve dispenser, crosslinked with the aerosols of CaCl_2_, and then incubated for two days to facilitate fusion amongst spheroids. As shown in **Figure 2D**, two days after bioprinting and incubation in alginate, the ∼1 mm pyramid topography, assembled using hMSC/HUVEC spheroids, maintained its defined shape with evidence of organized cytoskeletal filaments. After removal of alginate and further incubation, the constructs were observed to undergo significant compaction, lost their designed configuration and turned into tissue balls (**Figure S1**), which is corroborated by previously published articles [14,22,30]. In order to direct stem cell differentiation towards bone tissue formation, constructs composed of stem cells should be inducted for about three weeks [14,16,31]. In this case, the bioprinted hMSC/HUVEC construct deformed into a tissue aggregate within four days of being guided into bone tissue differentiation. Hence, the concern with inherent compaction of spheroids leading to shape contortion hinders the whole objective of bioprinting spheroids in complex desired geometries. This led us to explore another strategy, by tuning the timelines of induction of osteogenic differentiation, for retaining the geometry of constructs while bioprinting using spheroids.

### 3.3 Osteogenic differentiation of hMSC/HUVEC spheroids

To induce osteogenesis, hMSC/HUVEC spheroids were cultured for five days in growth media and the following 10 days in osteogenic media. The core of the spheroids was visualized by H&E staining of the cross-sections. As shown in **Figures 3A1 and 3B1**, hMSC/HUVEC spheroids, after induction to osteogenic lineage, showed more compact and condensed structures with uniformly distributed cells compared to that in hMSC-only spheroids. Mainly, this phenomenon is more pronounced at the center of spheroids, with ring like structures at the periphery of the spheroids. This is due to the presence of HUVECs in cellular spheroids [16]. Osteogenic cells begin the deposition of mineral deposits when they differentiate into osteoblasts [32]. Calcium deposition of hMSC-only and hMSC/HUVEC spheroids was confirmed by alizarin red S (ARS) staining (**Figures 3A2 and 3B2**). The deposited calcium amount of hMSC/HUVEC spheroid was substantially higher than that of hMSC-only spheroid with uniform calcium deposition across the entire cross-section, except for the periphery of the spheroid. The presence of the ring like structure (H&E) and absence of calcium deposition (ARS) at the periphery of the hMSC/HUVEC spheroids led us to characterize the localization of the hMSCs and HUVECs in the spheroids after osteogenic induction. RUNX2, an early osteogenic marker, was verified by immunocytochemical staining along with CD31 staining to confirm the cells of endothelial lineage. RUNX2 marker in hMSC/HUVEC spheroids was uniformly distributed across the cross-section of the spheroid while lesser expression of RUNX2 was observed in hMSC-only spheroid (**Figures 3A3 and 3B3**). Besides, hMSC/HUVEC spheroids showed more endotheliogenic-specific marker than hMSC-only spheroids, as indicated by positive staining of CD31, with more localized CD31+ cells at the periphery of the spheroid, forming a ring like structure, which is in agreement with previous findings [33]. Thus, hMSC/HUVEC spheroids demonstrated enhanced proliferation and increased mineralization which is direct evidence of enhanced osteogenic potential due to a synergistic display between the two cell types [34,35]. Crosstalk between hMSCs and endothelial cells is not only dependent on microenvironmental diffusion factors but also, on gap junction alpha-1 protein [36] which demonstrates the necessity of direct cell-cell contact between hMSCs and HUVECs [37]. Besides this juxtacrine mechanisms by the gap junctional activity, endothelial cells mediated paracrine factors including vascular endothelial growth factor (VGEF) and the inflammatory mediator prostaglandin E_2_ (PGE2) augments the crosstalk between periodontal ligament stem cells (PDLSCs) and endothelial cells. This, in turn, enhances the osteogenic potential of PDLSCs under hypoxia regulated by mitogen-activated protein kinase (MAPK) kinase/extracellular signal–regulated kinase and p38 MAPK pathways [18]. As the core of the spheroids are deemed more hypoxic compared to the periphery, osteogenic differentiation of stem cells is enhanced by the PGE_2_ and VEGF factors secreted by the HUVECs. HUVECs have also shown to express bone morphogenic protein 2, which induces osteogenic differentiation of stromal stem cells [38]

**Figure 3.**
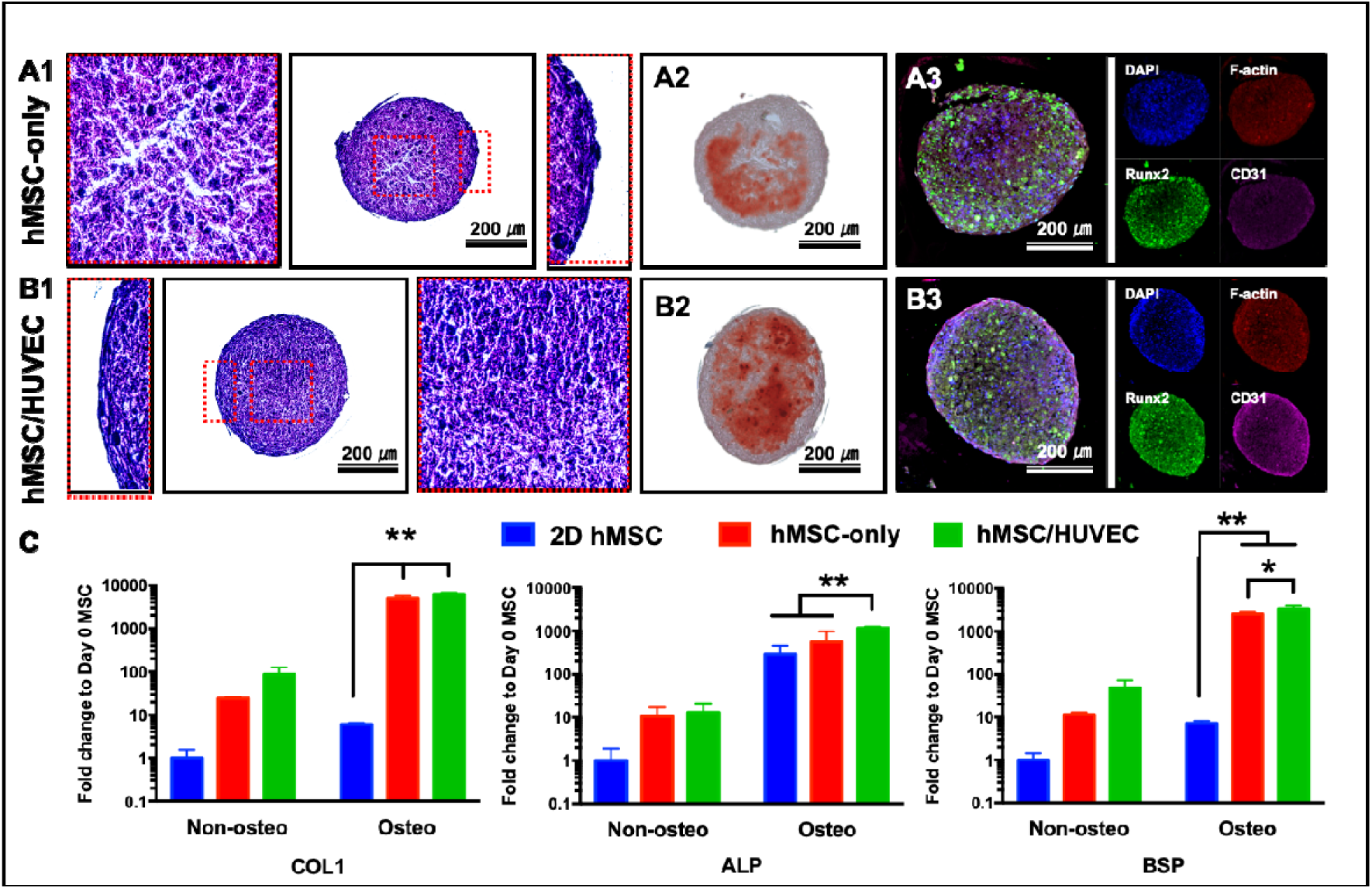
Characterization of osteogenically differentiated hMSC/HUVEC spheroids, cultured for five days in growth media and 10 days in osteogenic media: Representative images of **(A1-B1)** H&E staining, **(A2-B2)** ARS staining, and **(A3-B3)** IF staining with RUNX2 (Green), F-actin (Red), CD31 (Magenta), and cell nuclei (Blue) of hMSC-only and hMSC/HUVEC spheroids respectively. **(C)** Osteogenic related gene expression levels of 2D hMSC, hMSC-only, and hMSC/HUVEC spheroids cultured in growth and osteogenic media; *p < 0.05 and **p < 0.01. The results demonstrate the efficacy of HUVECs incorporation in hMSC spheroids to enhance osteogenic differentiation across the entire spheroid domain.

With considerable qualitative evidence demonstrating the effects of HUVECs on osteogenic potential of hMSCs, we also analyzed the mRNA expression of type I collagen (COL1), alkaline phosphatase (ALP), and bone sialoprotein (BSP) to quantify the same. All the experimental groups consisting of 2D hMSC, hMSC-only spheroids and hMSC/HUVEC spheroids were divided into two categories − cultured in either the maintenance media (labeled as “Non-osteo”) or osteogenic media (labeled as “Osteo”). As shown in **Figure 3C**, all groups cultured in osteogenic media exhibited higher expression of COL1, ALP, and BSP than those in maintenance media. The early-stage osteogenic differentiation markers of COL1 and ALP increased in the order of 2D hMSC, hMSC-only, and hMSC/HUVEC in both types of culture media. COL1 expression for hMSC-only and hMSC/HUVEC spheroids was significantly higher than that for 2D hMSC. This demonstrates the efficacy of 3D spheroids to accurately demonstrate the influence of microenvironmental factors on osteogenic differentiation ^22^. Also, hMSC/HUVEC spheroids cultured in osteogenic media exhibited significantly increased ALP expression compared to both 2D hMSC and hMSC-only spheroids cultured similarly. This is a direct evidence that the endothelial cells enhance the differentiation of hMSCs. Although, the precise role of HUVECs on osteogenic differentiation is debated, many 2D co-cultures with hMSCs and HUVECs have reported such observations [39,40]. The expression of BSP, a late osteogenic differentiation marker, significantly increased in the order of 2D hMSC, hMSC-only, and hMSC/HUVEC spheroids similar to the COL1 and ALP results. In particular, 3D hMSC/HUVEC showed the highest expression of BSP compared to other groups. Overall, it was found that hMSC/HUVEC spheroids exhibited enhanced early and late stage osteogenic differentiation, both in presence and absence of osteogenic media corroborating our earlier H&E, ARS and immunocytochemical staining and previous literature [39,40].

Therefore, hMSC/HUVEC spheroids were used for bioprinting of osteogenic tissues. Quiescence is inherently associated with niche resident stem cells. Previously studies have reported lower hMSCs proliferation rates in a co-culture system with HUVECs [33], suggesting the role of HUVECs in a retardation of hMSCs activity. Inducing the hMSC/HUVEC spheroids to differentiation media before bioprinting lowers the proliferative potential of hMSCs and allow the spheroids to be in their osteogenic differentiation pathway [41]. Hence, in this study, we utilized mid-term osteogenic hMSC/HUVEC spheroids for bioprinting scaffold-free bone-tissue constructs with an expectation of reduced deformation in the bioprinted geometry.

### 3.4 3D bioprinting of osteogenic tissues

To demonstrate the ability of osteogenic hMSC/HUVEC spheroids to serve as building blocks in bioprinting, different topographies resembling triangle, hexagon, and diamond in a single layer were bioprinted using the AAB system. HMSC/HUVEC spheroids cultured for five days in maintenance media and differentiated for 10 days in osteogenic media were bioprinted on the gel substrate, overlaid with the sacrificial alginate, and then incubated for another two days in the osteogenic media. As shown in **Figure 4A**, hMSC/HUVEC spheroids were precisely arranged into a diamond shape and after incubation, fused into a single patch of tissue. We also demonstrated a diamond shaped topography in three layers with osteogenic hMSC/HUVEC spheroids (**Figure 4B**). Bioprinted single-layered osteogenic tissue construct exhibited high cell viability from the periphery to the core of the spheroid with a negligible number of dead cells, as indicated by LIVE/DEAD staining. In histological analysis, H&E-stained tissue sections revealed that osteogenic hMSC/HUVEC spheroids were tightly self-assembled to each other, and bioprinted tissues maintained their original shape without falling apart. The histological analysis of the cross-sections of the 3D diamond-shaped construct by H&E presented uniform cell distribution through the entire 3D structure with a very well-fused interface amongst the spheroids. Mineralization was confirmed by measuring mineralized volume and calibrated bone mineral density by μCT analysis. The results showed that the single-layered bioprinted constructs were predominantly composed of mineralized matrix, which was indicated by values exceeding the threshold range of 200 mgHA/ccm. The mineralized volume in the single-layered triangle, hexagon and diamond shaped configurations was measured to be 0.051, 0.056 and 0.081 mm^3^, respectively. Uniform mineralization was also observed in the 3-layered diamond topography, with no significant alternation in its shape. The mineralized volume in first, second- and third-layer of the 3-layered geometry was measured at 0.163, 0.036 and 0.004 mm^3^, respectively.

**Figure 4.**
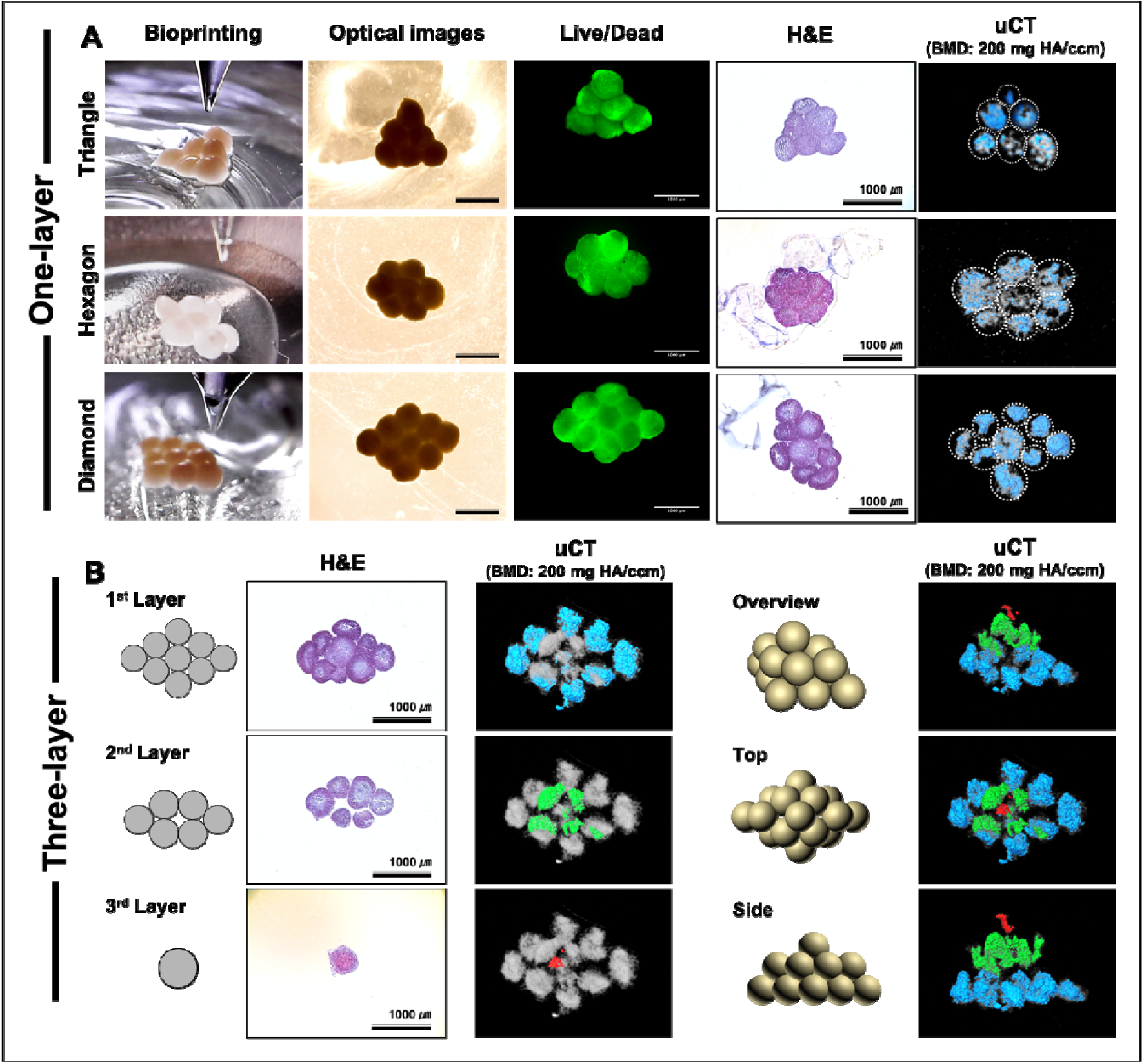
3D biofabrication of (A) triangle, hexagon, and diamond-shaped topographies in a single layer (A) and 3 layers (B) using mid-term osteogenic differentiated hMSC/HUVEC spheroids. The bioprinted tissue constructs were characterized by optical microscopy, LIVE/DEAD assay, H&E and μCT showing bone formation and mineralization two days post bioprinting. The red, green and blue colors in the overview uCT scans depict the 3^rd^, 2^nd^ and 1^st^ layer, respectively, of the 3-layered diamond-shaped bioprinted tissue. We show negligible shape changes in the 3D tissue constructs post-bioprinting utilizing pre-differentiated hMSC/HUVEC spheroids.

Thus, in this study, we presented a new approach to fabricating scaffold-free bone tissue constructs using the AAB system with mid-term osteogenic hMSC/HUVEC spheroids as building blocks. The constructs retained the bioprinted shape and demonstrated uniform mineralization throughout the entire 3D configuration.

This practical AAB approach could be refined in the near-future to enhance the throughput and scalability and has the likelihood to be used as a reproducible tissue bioprinting platform, for various applications, including organ-on-a-chip devices, microfluidics, organoid engineering and tissue engineering. In addition, since lack of vascularization is one of the major roadblocks for functional bone tissue constructs[42] leading to tissue necrosis, long-term bone tissue viability has to be addressed in the future by the fabrication of scaffold-free constructs with an extensive and efficient network of vascular micro-vessels is desirable, prior to preclinical testing.

## 4. Conclusion

In conclusion, we present an effective spheroid bioprinting strategy using the AAB system, which facilitates bioprinting of osteogenic hMSC/HUVEC spheroids to fabricate scaffold-free 3D bone tissue constructs. This study addresses the limitations of the previously explored spheroid bioprinting techniques by enabling precise positioning of spheroids onto a hydrogel (11% with respect to the spheroid diameter), preservation of the geometrical shape of the fabricated construct by reducing compaction, and induction of uniform bone formation across the entire spheroid. HUVECs were introduced into hMSCs to investigate their influence on spheroid formation and compaction, osteogenic differentiation and reduction of shape deformation after differentiation. Our findings demonstrate that osteogenically-differentiated hMSC/HUVEC spheroids, with as little as 8% of HUVECs, could be used as building blocks for bone tissue fabrication. These spheroids demonstrated reduced necrosis, increased cell viability in the core of the spheroid, enhanced differentiation into osteogenic lineage, and improved mechanical properties. Additionally, our bioprinting strategy exploited the cohesive nature of spheroids and reduced the proliferative potential of hMSCs after differentiation; this enabled building a scaffold-free 3D construct without any major shape change after bioprinting. Such a bioprinting approach, to facilitate fabrication of 3D geometries with negligible shape changes, using spheroids composed of differentiated stem cells and introduced with endothelial cells to enhance cell viability and differentiation, provides a new direction in bottom-up, scaffold-free bone tissue fabrication.

## 5. Acknowledgement

This work has been supported by National Science Foundation Awards 1914885 and a Convergence grant from the Materials Research Institute at Penn State University. D.N. H. acknowledges the support from the National Research Foundation of Korea (NRF) grant funded by the Korean Government (MIST) (2017R1D1A1B04030398) (2020R1C1C1007129).

## Contributions

D.N.H., B.A. and I.T.O. designed the research. D.N.H., B.A., M.D., and H.W. performed the experiments, and D.N.H., B.A. D.B., G.L., and I.T.O. evaluated the results. All authors contributed to writing of the manuscript and agreed the final content of the manuscript.

## Data availability

All data needed to evaluate the conclusions in the paper are present in the paper and/or the Supplementary Materials. Additional data related to this paper may be requested from the authors.

## Ethics declarations

The authors declare no competing interests.

## Supplementary Video

**Supplementary Video 1**. 3D reconstruction of pyramid shape pattern

## Supplementary Information

**Figure S1.**
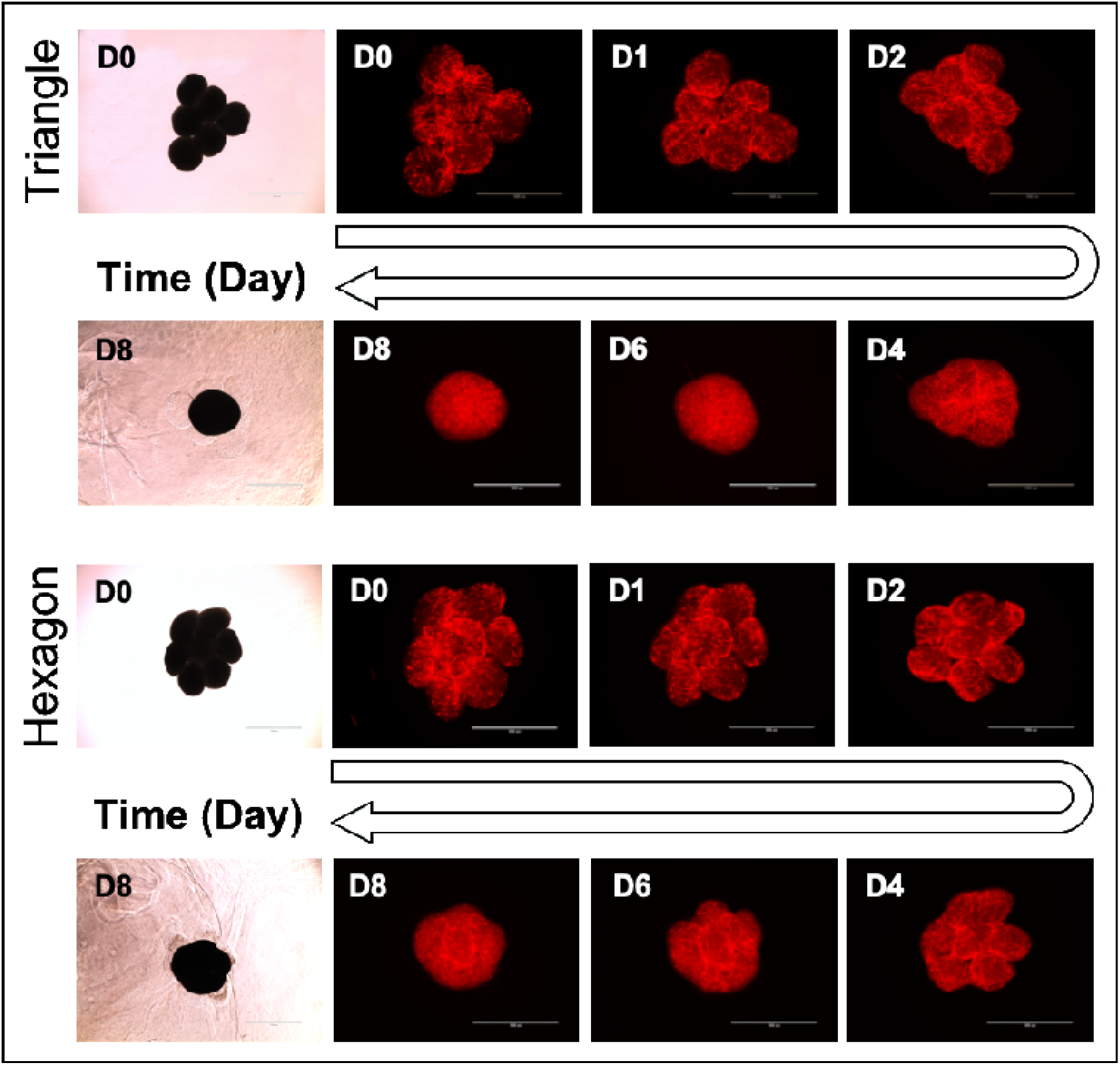
Post-bioprinting, hMSC/ tdTomato^+^ HUVEC tissue constructs were induced to osteogenic differentiation. The tissue constructs lost their shape and turned into tissue balls over a function of time (D: days).

